# Increased basal ganglia volume in older adults with tinnitus

**DOI:** 10.1101/2025.06.08.658513

**Authors:** Simón San Martin, Vicente Medel, Hayo Breinbauer, Carolina Delgado, Paul H. Delano

## Abstract

Tinnitus is the perception of sounds without external stimuli, affecting 10%-15% of the general population and up to 25% of individuals over 70 years of age. While traditionally viewed as an auditory phenomenon, growing evidence highlights the role of the central nervous system in its pathophysiology. One of the proposed mechanisms, the “gating hypothesis” of tinnitus, suggests an alteration in the modulation of sensory activity by the frontostriatal network. Although structural changes in frontal areas support this idea, gray matter differences in subcortical regions—such as the auditory pathway and basal ganglia— remain poorly understood. Here, we examined subcortical structures and auditory function in older adults with mild presbycusis from the ANDES cohort, including 51 tinnitus patients and 40 age-matched controls. We analyzed brain volume via structural magnetic resonance imaging and subcortical auditory functionality via auditory brainstem responses (ABRs). We found non-significant differences in age, hearing loss, cognitive performance, and ABR amplitudes between the groups. Notably, tinnitus patients presented a significant increase in the volume of basal ganglia structures (striatum and pallidum) but not in auditory areas. These findings reinforce the role of the basal ganglia in age-related tinnitus pathophysiology.

## Introduction

The perception of phantom sounds in the absence of external auditory stimuli, known as tinnitus, affects approximately 10% to 15% of the global population ^1–3^. Tinnitus can manifest solely as a perceptual symptom with no other coexisting alterations, but for a subset of 1% to 3%, it becomes a distressing syndrome linked to emotional and cognitive alterations, such as anxiety, insomnia and depressive symptoms ^4,5^. This difference has been identified as a key factor to consider in tinnitus research, highlighting the need to clearly define whether tinnitus corresponds to isolated tinnitus (sensory tinnitus) or to a tinnitus disorder with emotional and cognitive comorbidities ^6^. Importantly, the differential pathophysiological mechanisms or neuroplasticity changes underlying these two tinnitus phenotypes (sensorial tinnitus vs. tinnitus disorder) are not well understood ^7^. Elucidating the mechanisms of these two phenotypes of tinnitus can aid in the development of therapeutic strategies for alleviating the burden of tinnitus.

Several theoretical frameworks have been proposed to elucidate the neural mechanisms underlying tinnitus perception ^8–10^. Nearly 50% to 80% of tinnitus patients have peripheral hearing loss due to inner ear damage, such as the loss of cochlear hair cells produced by acoustic trauma, ototoxicity, or age-related hearing loss ^11–14^. The central gain hypothesis proposes that, owing to the diminished afferent activity in the cochlea and auditory nerve of individuals with hearing loss, there is an increase in neural activity within central auditory structures that could lead to tinnitus perception ^4,15,16^. However, recent evidence suggests that the increased central gain activity in tinnitus patients may be more closely related to the presence of hyperacusis (intolerance to loud sounds) than to tinnitus itself ^10,17^.

Another proposed mechanism in tinnitus involves an alteration in the top-down gating of auditory activity by frontostriatal and basal ganglia circuits ^18–21^. In this sense, dysfunction in sensory gating could contribute to the persistence and distress associated with tinnitus ^21–23^. Moreover, there are clinical reports showing that deep brain stimulation of the basal ganglia (caudate nucleus) can ameliorate tinnitus severity ^24^; consequently, this subcortical structure has been proposed as a target for the neuromodulatory treatment of tinnitus ^25,26^. However, the specific contribution of a basal ganglia gating mechanism in tinnitus in older adults is mostly unknown.

Aging and hearing loss are important factors to consider in older adults with tinnitus ^9,27–29^. The incidence of tinnitus is known to increase with age and hearing loss ^30^, affecting the quality of life of older adults ^30–32^. In addition, hearing loss and aging are also associated with structural and functional brain changes ^33–35^, making it difficult to determine structural brain changes that are specifically related to the presence of tinnitus in older adults ^36^. In addition, although it is well established that aging and hearing loss are risk factors for cognitive decline and dementia ^37,38^, whether tinnitus is associated with cognitive decline remains controversial ^39,40^.

Importantly, a variety of functional and anatomical changes in different brain regions have been reported in MRI studies in tinnitus patients ^33,36,41–44^. For example, previous studies have reported both volume reductions ^43^ and increases ^42^ in basal ganglia structures among tinnitus patients, making this a controversial issue. These differences could be explained by the clinical heterogeneity of tinnitus and its comorbidities, such as the presence/absence of hearing loss, the aging process or cognitive decline ^5,6^. In this sense, it is important to study tinnitus characteristics and pathophysiological mechanisms in a specific group of older adults with tinnitus.

Here, we propose investigating structural and functional brain changes in subcortical nuclei in a subset of older adults with normal hearing to mild presbycusis who report tinnitus and do not use hearing aids. We hypothesize that the frontostriatal and basal ganglia networks play key roles in the pathophysiological mechanisms of the perception of sensory tinnitus in older adults. To test this hypothesis, we examined brain structural characteristics via MRI and auditory brainstem responses (ABRs) via the Chilean ANDES cohort of older adults ^45,46^.

## Results

A total of 91 older adults from the ANDES cohort were included (55 females), with an average age of 73.4 ± 5.1 years (ranging from 65 to 85 years), a mean hearing threshold of 30 ± 13.2 dB HL (PTA 0.5-4 kHz), and a mean education level of 9.5 ± 4.1 years. We identified 51 subjects who reported having tinnitus (T) and 40 subjects who were classified as non-tinnitus (NT). There were no significant differences in age or sex distribution between the tinnitus and non-tinnitus groups, whereas we detected a significant difference in educational level, which was lower in the tinnitus group (T: 8.7 ± 4.4 years of education; NT: 10.6 ± 3.6 years of education, Figure 1A, p<0.04). A summary of the demographic, audiological, cognitive, and emotional variables is presented in Table 1.

**Figure 1.**
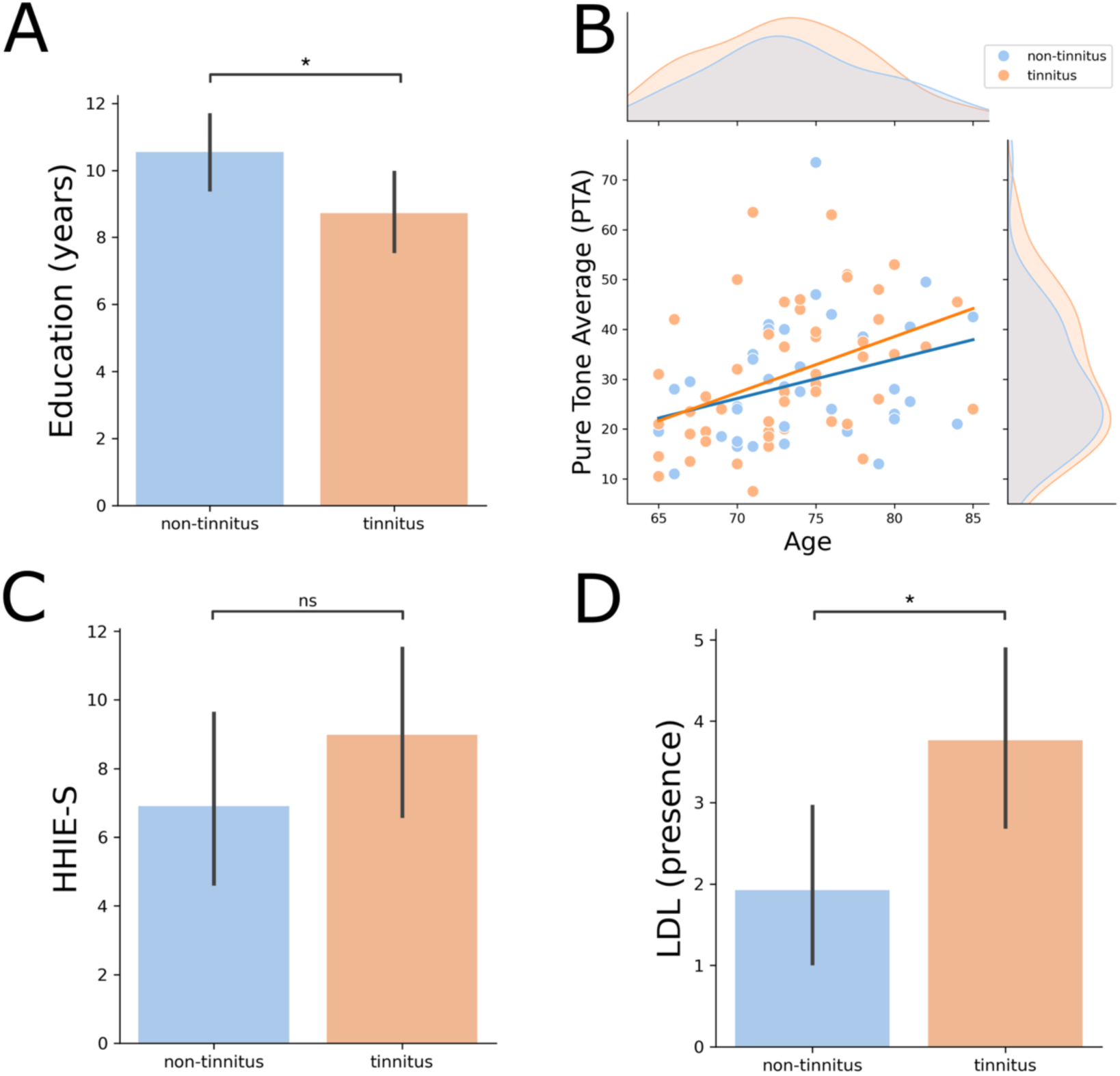
Educational level and audiological assessments. A) There were significant differences between groups in terms of years of education, being lower in tinnitus. B) The mean age and hearing levels of the tinnitus and non-tinnitus groups were similar. Significant correlations between PTA and age were found for both groups (T: Spearman rho=0.48, p=0.0003; NT: Spearman rho=0.38, p=0.013). C) The Hearing Handicap Inventory (HHIE-S) did not differ between the groups in terms of quality of life or emotional well-being with respect to hearing difficulties. D) Significant differences between groups were found in the loudness discomfort level (LDL), as the tinnitus group had a greater number of audiogram frequencies with detectable LDL, denoting a higher index for hyperacusis (T: 3.76 ± 4.1; NT: 1.92 ± 3.22, p=0.02). * = statistically significant, ns = non-significant.

**Table 1.** Demographic, audiological and cognitive variables of the two groups (tinnitus v/s non-tinnitus). * = statistically significant.

| Variable | Non-tinnitus (n = 40) | Tinnitus (n = 51) | Statistical comparison |
| --- | --- | --- | --- |
| Age, mean (SD) | 73.8 ± 5.2 | 73.1 ± 5.1 | t = 0.6 ; p = 0.54 |
| Female n (%) | 24(60) | 33(64.7) | X<sup>2</sup> = 0.06 ; p = 0.81 |
| Education, mean years (SD) | 10.6 ± 3.6 | 8.7 ± 4.4 | t = 2.1 ; p = 0.04* |
| MMSE, mean (SD) | 28.1 ± 1.32 | 28.0 ± 1.33 | t = 0.31 ; p = 0.75 |
| GDS, mean (SD) | 2.6 ± 2.4 | 3.5 ± 3.6 | t = -1.2 ; p = 0.21 |
| HHIE-S, mean (SD) | 6.9 ± 7.5 | 8.9 ± 9.1 | t = -1.5 ; p = 0.25 |
| PTA, mean (SD) | 29.1 ± 12.7 | 30.8 ± 13.8 | t = 0.59 ; p = 0.55 |
| LDL, count (SD) | 1.92 ± 3.22 | 3.76 ± 4.1 | U = 757.0 ; p = 0.02* |
| Wave I, mean (SD) | 0.127 ± 0.063 | 0.112 ± 0.068 | t = 1.04 ; p = 0.30 |
| Wave V, mean (SD) | 0.349 ± 0.111 | 0.377 ± 0.139 | t = 1.01 ; p = 0.31 |
| Ratio V/I mean (SD) | 3.85 ± 3.53 | 5.03 ± 3.99 | t = 1.46 ; p = 0.14 |

The presence of tinnitus disorder is usually accompanied by cognitive and emotional distress, whereas sensorial tinnitus is an isolated perceptual phenomenon. Using audiological, cognitive and mood surveys available in the ANDES cohort, we compared the tinnitus and non-tinnitus groups to determine the level of comorbidities in the tinnitus groups. We found non-significant differences in audiometric hearing levels (T: 30.8 ± 13.8 dB HL; NT: 29.1 ± 12.7 dB HL, p=0.55) and HHIE-S scores (T: 8.9 ± 9.1; NT:6.9 ± 7.5, p=0.25) between the tinnitus and non-tinnitus groups (Figures 1B and 1C). On the other hand, we found that the tinnitus group had greater audiogram frequencies with detectable LDL than the non-tinnitus group did (T: 3.76 ± 4.1; NT: 1.92 ± 3.22, p=0.02), indicating a greater presence of hyperacusis in the tinnitus group (Figure 1D). There were no significant differences in cognitive performance, as evaluated by the MMSE (T: 28.0 ± 1.33; NT: 28.1 ± 1.32, p=0.75), or depressive symptoms, as assessed with the GDS (T: 3.5 ± 3.6; NT: 2.6 ± 2.4, p=0.21), between the groups. Taken together, these results show that besides LDL and education, there were no significant differences in age, sex, hearing thresholds, MMSE scores or depressive symptoms between the groups with and without tinnitus in the ANDES cohort. We studied suprathreshold ABR responses to evaluate possible compensatory functional gains in the central auditory pathways of tinnitus patients ^16^. We found non-significant differences in the amplitudes of wave I (t=1.04, p=0.3), wave V (t=1.01, p=0.31) and the V/I ratio (t=1.46, p=0.14) between the tinnitus and non-tinnitus groups (Figure 2).

**Figure 2.**
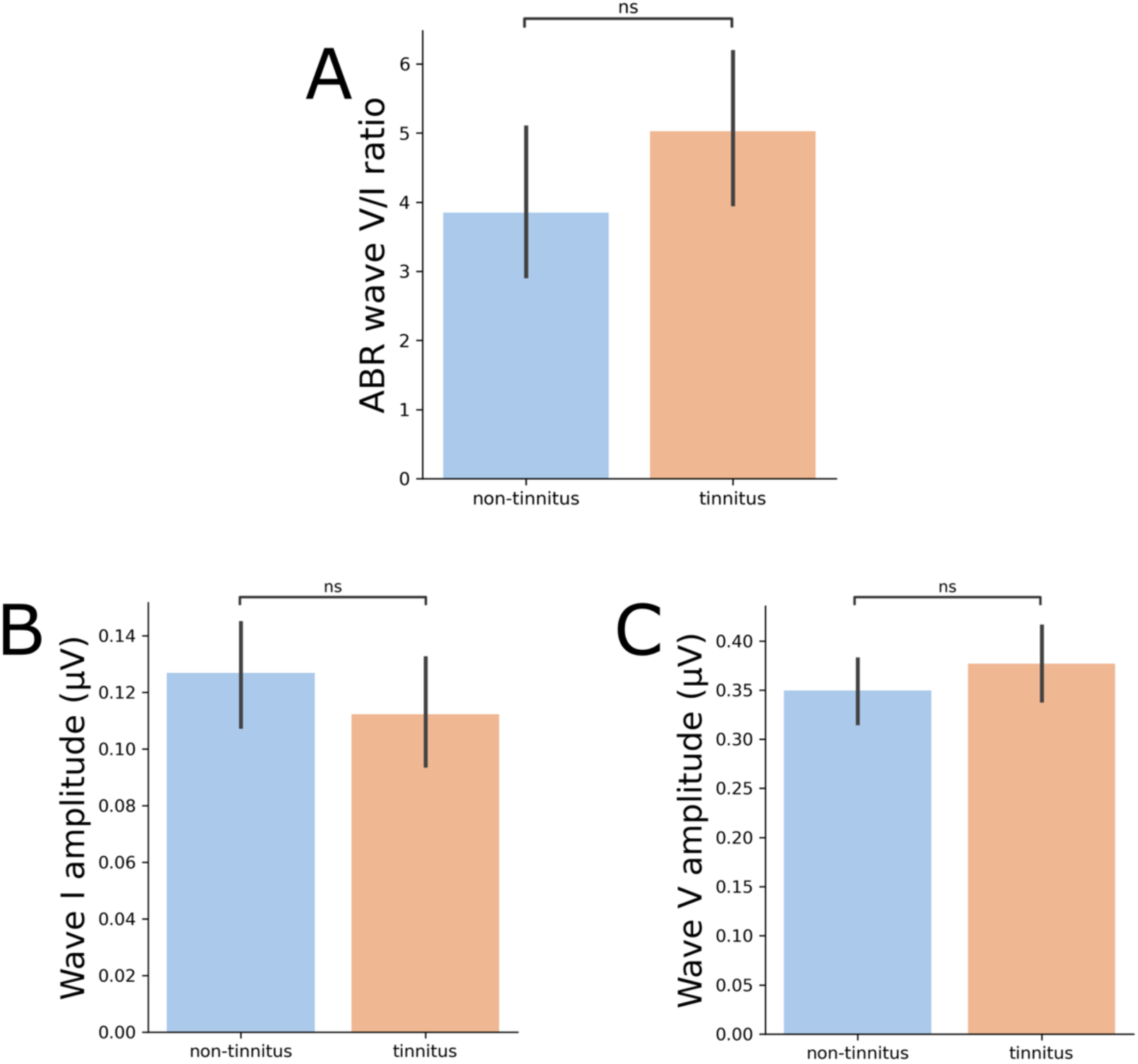
Functional evaluation of the central auditory pathway using suprathreshold ABR in the groups with and without tinnitus. A) The wave V/I ratio comparison revealed no difference between healthy controls and tinnitus patients. B) Non-significant difference in auditory-nerve (Wave I) amplitude. C) Non-significant difference in wave V amplitude. ns=non-significant.

Next, we used volumetric MRI to evaluate possible structural changes in the central auditory pathway and in subcortical non-auditory brain structures of both groups. First, we calculated the volume of the nuclei of the central auditory pathway, including the cochlear nuclei, superior olivary complex, inferior colliculi, and the medial geniculate body and auditory cortex, by using an atlas of the human auditory pathway developed at 7-Tesla MRI^47^. There were no significant differences between the tinnitus and non-tinnitus groups in the intracranial volume (t=-0.75, p=0.54), cochlear nuclei (t=-0.55, p=0.57), superior olivary complex (t=0.054, p=0.93), inferior colliculi (t=-0.2, p=0.81), or auditory cortex (t=-1.15, p=0.25) (Figure 3).

**Figure 3.**
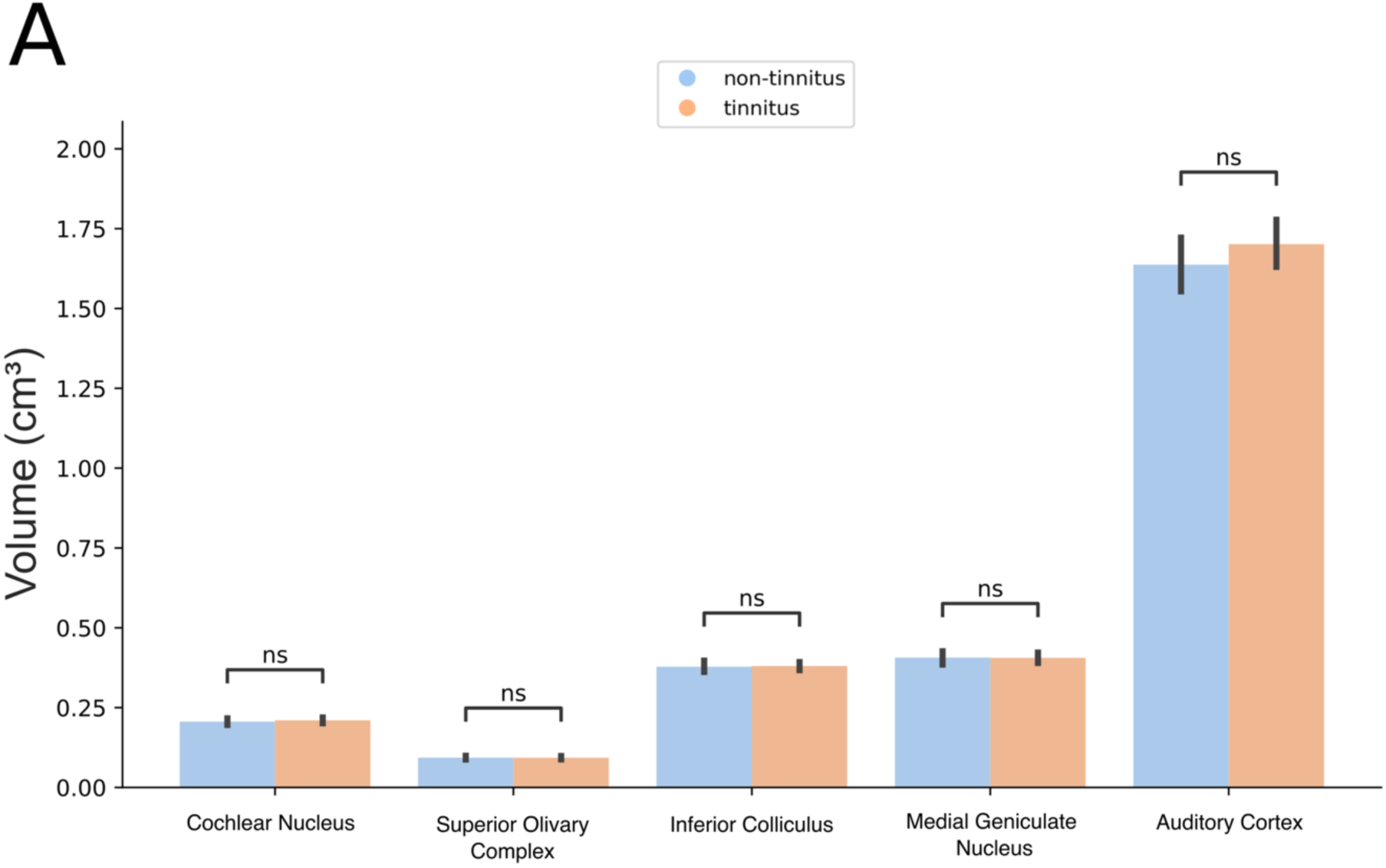
Volumetric MRI comparisons of the central auditory pathway in the groups with and without tinnitus. There were no significant differences between the groups in any of the structures measured. ns=non-significant.

Second, we evaluated structural changes in non-auditory brain subcortical structures, which are related to top-down gating mechanisms that could be altered in tinnitus patients ^15,23^. To test this hypothesis, we calculated the gray matter volume of these subcortical nuclei and compared them between groups. We found that for basal ganglia structures, the tinnitus group had significantly larger volumes (pallidum, t=-2.1, p=0.03; Figure 4A; putamen, t=-3.6, p=0.0004; Figure 4B; caudate, t=-2,8, p=0.0038; Figure 4C; accumbens, t=-3.4, p=0.0008; Figure 4D). Figure 5 shows the anatomical reference for the striatum structures analyzed and their corresponding t values calculated for the comparisons between groups ^48^. We then calculated the striatum volume by summing the putamen, caudate, and accumbens volumes and found a significant difference in striatum volume between groups, even after controlling for the estimated total intracranial volume and age of the subjects (ANCOVA striatum, 𝐹=24.36, 𝑝 value<0.001, 𝜂𝑝^!^=0.45).

**Figure 4.**
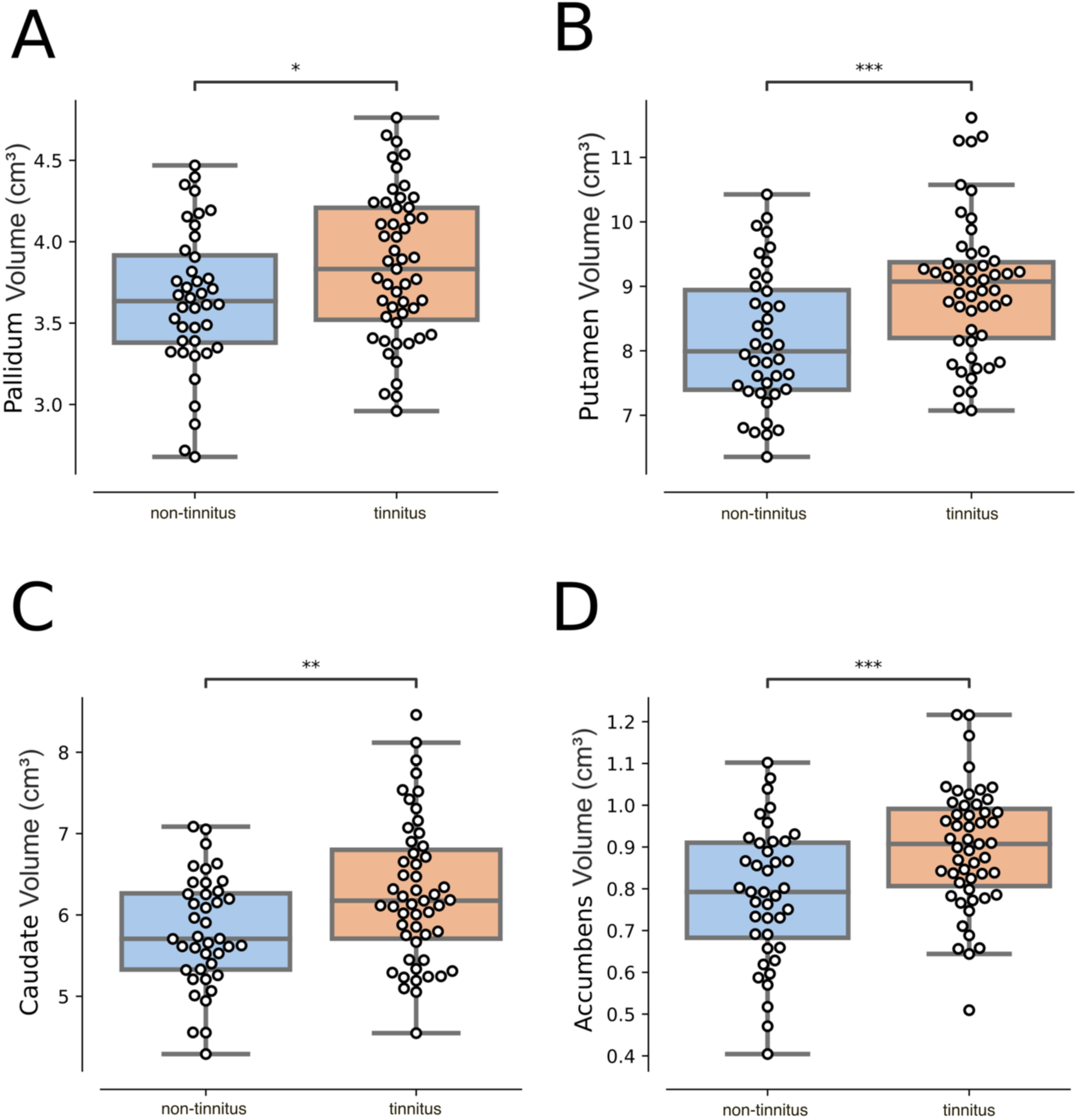
Basal ganglia volume comparisons. Volumetric comparisons of nonauditory subcortical structures revealed that tinnitus patients have larger pallidum (A), putamen (B), caudate (C), and nucleus accumbens (D) structures. *p<0.05, **p<0.01, ***p<0.001.

**Figure 5.**
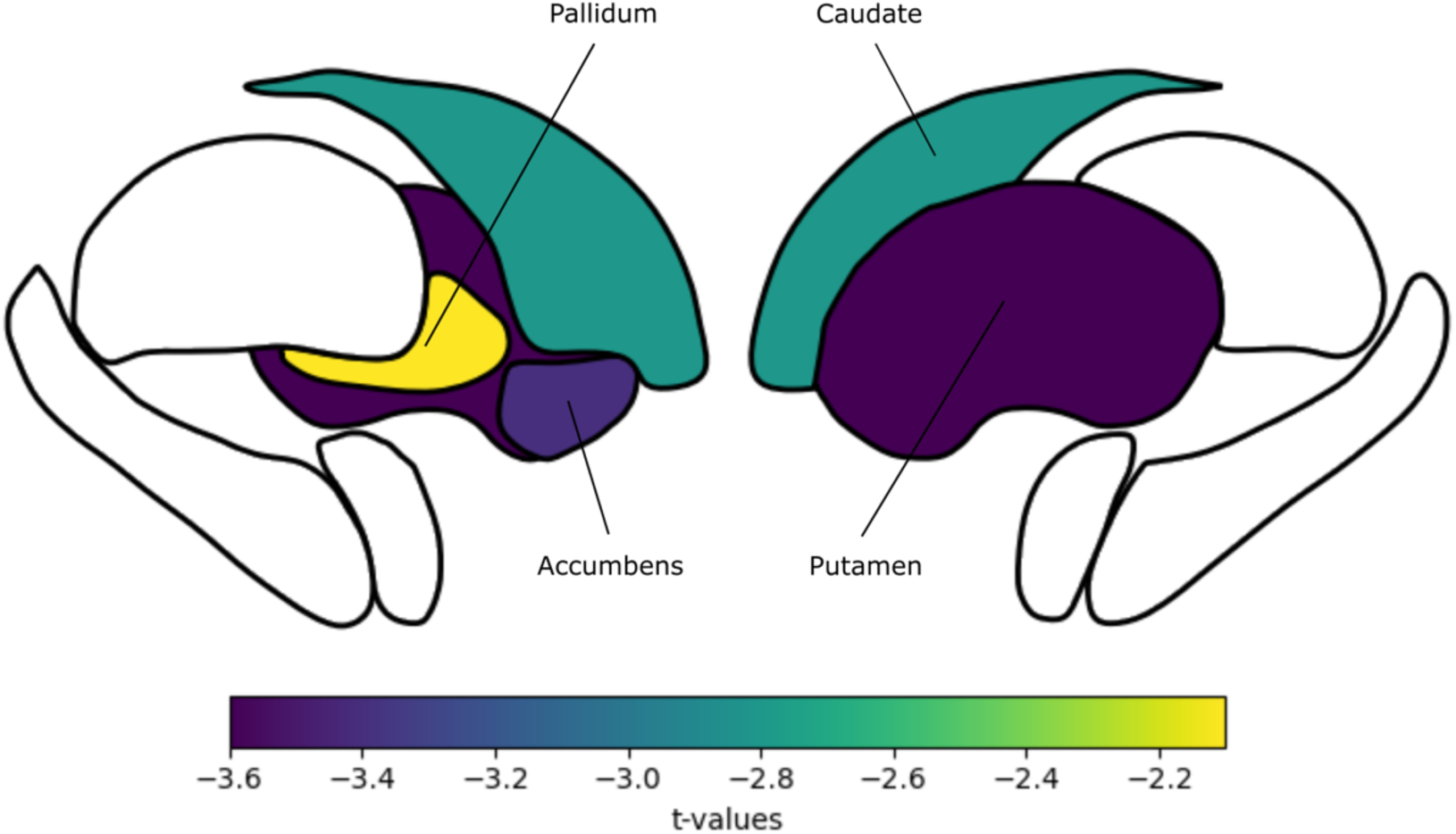
Anatomical reference for the basal ganglia structures analyzed, along with their corresponding t-values calculated for the statistical comparisons.

Afterwards, we conducted two robust linear regression models to examine whether the effect of tinnitus on basal ganglia volume was influenced by (i) age and (ii) hearing threshold (PTA). The interaction effect between tinnitus and age was statistically significant (β=0.1451, p=0.047), suggesting that the relationship between tinnitus and basal ganglia volume varies with age. However, the interaction effect between tinnitus and the PTA did not reach statistical significance (B=0.0407, p=0.186), indicating that the PTA does not strongly moderate this relationship (Figure 6).

**Figure 6.**
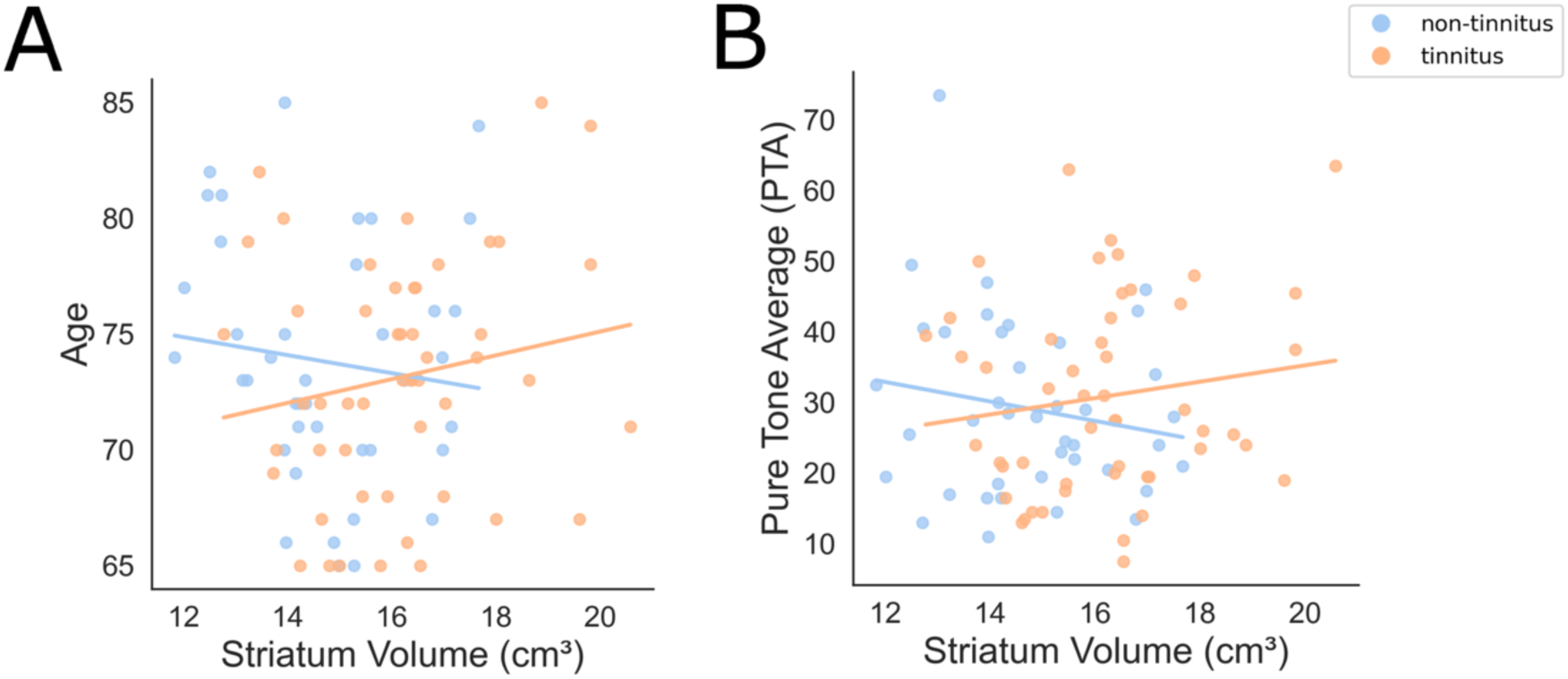
Interactions and effect of group (tinnitus vs. non-tinnitus) on striatum volume, with age (A) and PTA (B) as predictors. The striatum volume is the sum of the putamen, caudate, and accumbens volume structures.

## Discussion

The present results show that older adults who reported having tinnitus in the past year have lower levels of education and a worse hyperacusis index (LDL) compared to age- and hearing-matched non-tinnitus individuals. There were no significant differences in cognition or depressive symptoms between the groups. Additionally, we found non-significant differences between groups in the structure and function of the central auditory pathway. However, we observed increased volumes in the basal ganglia structures, including the striatum and pallidum, of tinnitus patients. We propose that individuals with tinnitus in the ANDES cohort primarily experienced sensory tinnitus, as they did not actively complain about it but were instead asked about its presence during audiometric evaluations.

### Sensorial tinnitus versus tinnitus disorder

Tinnitus, far from being a single-stereotyped disorder, encompasses a wide array of maladaptive neural network configurations for sound processing ^9,49^, ranging from sensorial or isolated tinnitus to tinnitus disorders with behavioral and emotional comorbidities ^6^. In this study, we analyzed a specific group of tinnitus patients, which likely corresponded to sensorial tinnitus in older adults with normal to mild hearing loss and non-significant differences in age, hearing loss, cognition and depressive symptoms from non-tinnitus-matched individuals. The educational level was worse in tinnitus patients, which could be explained by the reported greater exposure to acoustic injuries that can lead to tinnitus, such as acoustic trauma or ototoxicity, in less educated populations ^50,51^.

### Lack of increased central auditory gain in tinnitus

In this cohort, neither the degree of deafferentation (measured by PTA) nor age itself was significantly associated with the likelihood of having tinnitus. Similarly, we did not find differences in electrophysiological (wave amplitudes in ABR) or morphological (size of key auditory nuclei in the brainstem) markers of central gain between the tinnitus and non-tinnitus groups. Our results are in agreement with reports showing a lack of compensatory central gain in tinnitus patients ^7,17^. Importantly, the tinnitus group did present significantly worse LDLs, potentially indicating a reduced dynamic range of the auditory system—a characteristic of hyperacusis, which several authors propose as a clinical marker of increased central gain ^7,10,17^. Another explanation for the absence of central auditory gain increase in the tinnitus group of the ANDES cohort might be consequence of the slow progression of hearing loss and tinnitus in presbycusis patients. This slow progression may influence specific outcomes through better adaptation, such as the apparent lack of structural or electrophysiological markers of increased central gain in older adults with tinnitus.

### Proposed mechanisms for the increased volume in the striatum and pallidum

In this specific cohort, the basal ganglia emerged as structures potentially linked to tinnitus. In recent decades, increasing evidence has shown that the basal ganglia play functional roles beyond movement control ^52^. Basal ganglia circuits constitute highly conserved vertebrate structures receiving connections from diverse brain sources related to sensorimotor, visual, auditory, premotor, and emotional processing, among others ^53,54^.

Dysfunctions in the basal ganglia have been implicated in various conditions, including motor disorders (e.g., Parkinson’s disease, Huntington’s disease), psychotic disorders (e.g., schizophrenia), Tourette syndrome, obsessive‒compulsive disorder, and addictive behaviors55.

Research on the brain structure of tinnitus patients has reported contradictory results concerning structural changes in the basal ganglia, including decreases in volume ^43^ and increases in volume ^42^. In our case, we are in agreement with investigations reporting basal ganglia volume increases, which we propose for this specific subset of tinnitus individuals: that is, older adults with sensory tinnitus. In this sense, the results obtained in the group with tinnitus could be representative of beneficial neural adaptations to tinnitus in older adults without cognitive or depressive comorbidities.

The latter proposition can be supported by recent evidence obtained in patients treated with electroconvulsive therapy (ECT) for resistant depression. Brain MRI after ECT revealed gray matter volume increases in several regions, including the basal ganglia, suggesting that the increase in gray matter volume might be related to the clinical improvement of depressive symptoms ^56,57^. Therefore, we propose that the increased volume changes that we found in the basal ganglia of the tinnitus group might have contributed to maintaining the phenotype of sensory tinnitus and preventing the emergence of tinnitus disorder syndrome. These findings could support the hypothesis that the basal ganglia may play a role in the inhibition attempt to suppress tinnitus-related activity within auditory circuits rather than generating the tinnitus signal itself ^58^. A speculative scenario could be that, in older adults with sensorial tinnitus, inappropriate signals generated in the auditory pathway are filtered by striatal circuits. This filtering interrupts a vicious cycle that might otherwise establish these changes as more permanent, preventing the emergence of tinnitus syndrome with cognitive and depressive comorbidities. However, the literature and our study (cross-sectional) do not allow for confirmation of a causal relationship between striatal function and top-down gating of tinnitus activity ^20,58^.

## Limitations

Among the limitations of this study, it is important to note that the primary methodology of the ANDES study did not focus specifically on tinnitus, nor did it include standard tinnitus intensity measures (e.g., tinnitus handicap inventory), which could provide insight into tinnitus severity and quality-of-life impacts. Future studies in this area would benefit from incorporating these parameters to explore potential correlations between basal ganglia volume and tinnitus severity in older adults.

## Conclusion

In this specific group of older individuals with tinnitus, the only significant difference in brain structure between the tinnitus group and the non-tinnitus group was the volume enlargement of the basal ganglia nuclei. There were no significant differences in subcortical auditory volume, ABR response, cognition or depressive symptoms between the two groups. These results suggest that plastic changes in frontostriatal networks may be relevant for avoiding the cognitive and emotional comorbidities that constitute tinnitus syndrome. Future work could identify basal ganglia networks as potential targets for tinnitus mitigation.

## Methods

### Participants

Ninety-one older adults from the Auditory and Dementia Study (ANDES) cohort were recruited from Recoleta’s primary public health centers in Santiago de Chile. Importantly, these patients did not complain about tinnitus, as they were recruited from primary care adult health screening programs, representing a specific subset of “sensorial tinnitus” in older adults. The inclusion and exclusion criteria were as follows: (i) were over 65 years of age at the time of recruitment, (ii) had no history of neurological or psychiatric illness, (iii) had no cause of hearing loss other than presbycusis (e.g., conductive hearing loss), and (iv) did not use hearing aids. The Ethics Committee of the Clinical Hospital of the University of Chile approved all procedures (permission number OAIC752/15). The participants provided written informed consent in accordance with the Declaration of Helsinki.

### Audiological assessment

Evaluations were carried out in the Department of Otolaryngology at the Clinical Hospital of the University of Chile. Audiometric hearing thresholds were evaluated from 0.125 to 8 kHz for each subject in both ears via a clinical audiometer (AC40, Interacoustics®). Conductive hearing loss was ruled out for each subject. The pure tone average (PTA) at 0.5, 1, 2, and 4 kHz was calculated for each subject in both ears.

Hyperacusis was estimated in dB (HL) using the loudness discomfort level (LDL) measured with an audiometer between 0.25 and 4 kHz. Owing to the frequent absence of LDL even at the maximum audiometer output (110 dB at 0.25 kHz and 115 dB between 0.5 and 4 kHz), group analyses for this variable focused on the detection or absence of a discomfort threshold at six frequencies (0.25, 0.5, 1, 2, 3 and 4 kHz) per ear, totaling twelve frequencies per subject. These analyses allowed us to use the whole sample of individuals, including those with no LDL, which have “0”, compared with subjects with LDL at all the evaluated frequencies, which yielded “12” summing both ears. Greater LDL values indicate more audiogram frequencies with hyperacusis.

Auditory brainstem response (ABR) waveforms were obtained by averaging with alternating clicks presented at suprathreshold levels (2000 repetitions, 80 dB nHL, bandpass 0.1–3 kHz, stimulus rate 21.1 Hz, EP25, Eclipse, Interacoustics®). The amplitudes of waves V and I were measured from peak to valley, and the latency times were calculated from the peaks of these waves. For the calculation of the V/I wave ratios, in situations where wave I was not detectable, the smallest amplitude value for wave I (0.02 μV) was utilized, as reported previously ^59^.

### Tinnitus

The audiologist of the ANDES cohort asked each participant during the audiometric evaluation about the presence or absence of tinnitus in the last year. The participants were divided into two categories: individuals experiencing tinnitus and those without tinnitus (non-tinnitus). The ANDES cohort does not include tinnitus intensity surveys.

### Image acquisition

Imaging data collection was performed via a MAGNETOM Skyra 3 Tesla full-body MRI scanner (Siemens Healthcare GmbH, Erlangen, Germany), which employs a T1 MPRAGE sequence. We captured continuous images of the entire brain via specific parameters for a shorter acquisition time: time echo (TE) at 232 ms, time repetition (TR) at 2,300 ms, and a flip angle of 8 degrees, consisting of 26 slices, with a matrix size of 256 × 256 and a voxel dimension of 0.94 × 0.94 × 0.9 mm³. Additionally, T2-weighted turbo spin echo (TSE) imaging (4,500 TR ms, 92 TE ms) and fluid-attenuated inversion recovery (FLAIR) imaging (8,000 TR ms, 94 TE ms, 2,500 TI ms) were performed to evaluate structural irregularities. The total imaging process took approximately 30 minutes, resulting in 440 images per participant.

### Image preprocessing and analysis

We employed two different morphometric approaches. First, we performed preprocessing with FreeSurfer, version 6, running under Centos 6. A single Linux workstation was used for the T1-weighted image analysis of individual subjects as suggested by previous research ^60^. FreeSurfer processes cortical reconstruction ^61^ through several steps: volume registration with the Talairach atlas, bias field correction, initial volumetric labeling, nonlinear alignment to the Talairach space, and final volume labeling. The automatic “recon-all” function produces representations of the cortical surfaces. It uses both intensity and continuity information from the entire three-dimensional MR volume in segmentation and deformation procedures. It creates gross brain volume extents for larger-scale regions (i.e., the total number of voxels per region): total gray and white matter, subcortical gray matter, brain mask volume, and estimated total intracranial volume. The reliability between manual tracing and automatic volume measurements has been validated. The correspondence between manual tracings and automatically obtained segmentations was similar to the agreement between manual tracings ^62^. All the volumes were visually inspected, and if needed, the volumes were edited by a trained researcher according to standard processes. With this approach, we calculated cortical and striatal volumes. Second, to analyze specific auditory pathway structures, we used a previously validated MNI mask ^47^, which was created to filter central auditory pathway structures (Cochlear Nuclei, Superior Olivary Complex, Inferior Colliculi, and Medial Geniculate Nucleus). To use the mask, we preprocessed each T1-weighted image in SPM12 following the Voxel-Based Morphometry pipeline (VBM). Briefly, each T1-weighted image was tissue-segmented, followed by a study-specific DARTEL template where each image was iteratively aligned with DARTEL. To quantify the auditory regions of interest, only the resulting gray matter images were used.

### Clinical assessment

To obtain clinical information about general cognition, the presence of depression, hearing impairments, and general functionality, multiple scales were used. The subject’s general cognitive function was evaluated via the Mini-Mental State Examination (MMSE), a widely recognized and validated tool for assessing cognitive status ^63^. Each participant self-assessed depression via the Geriatric Depression Scale (GDS) ^64^. This test consists of 15 quick questions about various aspects of mental health and mood. Higher scores on the GDS indicate a greater likelihood of depression. The Hearing Handicap Inventory for the Elderly Screening (HHIE-S) was used as a self-assessment tool specifically designed to evaluate the impact of hearing loss on the quality of life of older adults ^65^. It consists of 10 questions that focus on how hearing difficulties affect an individual’s daily life and emotional well-being. Higher scores indicate a greater impact of hearing loss on an individual’s life and well-being.

### Statistical analysis

All the statistical analyses and graphs were generated via Python. Group differences between tinnitus controls and non-tinnitus controls were examined via independent-sample parametric and nonparametric tests for audiological, demographic, clinical and structural variables. Spearman rank correlation tests were used to evaluate relationships between variables such as pure tone average and age. Correlations were calculated separately for the tinnitus and non-tinnitus control groups to explore potential differences in associations. Robust linear regression models were employed to investigate interactions between group, pure tone average, and age, assessing their combined effects on auditory and structural brain measures. Finally, analysis of covariance (ANCOVA) was used to control for confounders such as total intracranial volume and age in comparisons of gray matter volume across the auditory and striatal regions, which ensured that any observed group differences were independent of these covariates. The results are reported with group values or statistics and p values, with a significance threshold set at p < 0.05.

## Author contributions

C.D. and P.H.D. conceived the study design and oversaw cohort recruitment. S.S.M., V.M., H.B., C.D., and P.H.D. collected the experimental data and edited the manuscript. S.S.M., V.M., H.B., C.D., and P.H.D. conducted the statistical analyses and reported the results. S.S.M., V.M., H.B., C.D., and P.H.D. reviewed the manuscript. S.S.M., V.M., and P.H.D. Ensured that all the sections adhered to the journal guidelines.

## Data availability statement

Data from this article are available from the corresponding author upon reasonable request.

